# PNPLA3 I148M Reduces Hepatic Triacylglycerol Secretion and Mitigates Left Ventricular Diastolic Dysfunction in MASH Diet Mice

**DOI:** 10.1101/2025.04.24.650494

**Authors:** Andrew J. Butcko, Parsa Kamali, Nivedita Tiwari, Abir A. Rahman, Grace Teskey, Kehinde A. Adeshina, Nikhlesh K. Singh, James G. Granneman, Emilio P. Mottillo

**Affiliations:** Hypertension and Vascular Research Division, Department of Internal Medicine, Henry Ford Hospital, Detroit, MI, 48202; Department of Physiology, Wayne State University School of Medicine, Detroit, MI, USA 48202; Wayne State University School of Medicine, Detroit, MI; Department of Ophthalmology Visual and Anatomical Sciences, Wayne State University School of Medicine, Detroit, MI; Center for Molecular Medicine and Genetics, Wayne State University School of Medicine, Detroit, MI, USA 48202

**Keywords:** Lipids, Cardioprotection, Metabolism, Heart, Genetics

## Abstract

Cardiovascular disease (CVD) is a leading cause of mortality among individuals with metabolic dysfunction-associated steatotic liver disease (MASLD). Paradoxically, the strongest genetic risk factor for MASLD, PNPLA3-I148M, is associated with a reduced risk of CVD. The mechanisms for how PNPLA3-I148M causes MASLD and preserves cardiac health are not well understood. To further investigate cardioprotective effects of PNPLA3-I148M we expressed human WT-PNPLA3, PNPLA3-I148M or a GFP control in the livers of PNPLA3-/- mice. In agreement with prior investigations, mice expressing PNPLA3-I148M displayed greater hepatic neutral lipid accumulation on chow and a Metabolic Dysfunction-Associated Steatohepatitis (MASH) promoting diet. Changes in hepatosteatosis between the groups was not due to alterations in whole body metabolic parameters, although WT-PNPLA3 promoted greater glucose intolerance on a MASH diet. Echocardiography revealed that hearts from PNPLA3-WT mice had changes in left ventricular mass and thickness following 16-weeks of MASH, but not chow diet, which was not seen in the GFP or PNPLA3-I148M groups. Moreover, PNPLA3-WT and GFP mice had reduced E/A ratios (diastolic function), post MASH diet, which was not detected in PNPLA3-I148M mice. In addition, PNPLA3-I148M reduced hepatic secretion of TAGs under a MASH, but not chow diet. The expression of PNPLA3-I148M modified the liver lipidome, while minimal effects were observed in the heart. No differences in atherosclerotic plaque formation were observed following 24 weeks of MASH diet. These findings indicate that hepatic PNPLA3-I148M promotes hepatosteatosis yet protects against diet induced cardiac remodeling and diastolic dysfunction primarily through reductions in hepatic TAG secretion.

## INTRODUCTION

Metabolic dysfunction associated steatotic liver disease (MASLD) is the leading cause of chronic liver disease worldwide (1). Globally, MASLD has an estimated prevalence of 30% in the adult population, which is predicted to continue to rise (2). Development of MASLD is known to be multifactorial in its etiology with diet, genetics and environmental factors all playing a crucial role (3–5). Moreover, the pathogenesis of MASLD is largely thought to result from aberrant hepatic lipid metabolism which causes an imbalance of fat stores within the liver, leading to the accumulation of triacylglycerol (TAG) within hepatocytes (6). A common single nucleotide polymorphism in the patatin-like phospholipase domain-containing protein 3 (*PNPLA3*) gene (C>G: rs738409 mutation), resulting in an I148M amino acid substitution, is the greatest genetic determinant for development and progression of MASLD (7–10).

Despite its prominence, the exact mechanisms for how the I148M variant promotes MASLD is unclear. Interestingly, in murine models, initial studies demonstrated that neither whole-body deletion nor hepatic over-expression of wildtype (WT) PNPLA3 leads to MASLD, while overexpression of PNPLA3 I148M promotes TAG accumulation; observations which together suggest that PNPLA3 I148M is not a simple loss of function mutation, but acts in a dominant negative manner (11–14). More recent evidence suggests that the I148M variant promotes hepatosteatosis through both a reduction in PNPLA3’s intrinsic lipase activity, as well as its ability to sequester lipase co-activator alpha/beta hydrolase domain-containing protein 5 (ABHD5) away from phospholipase domain-containing protein 2 (PNPLA2), a closely related paralog of PNPLA3 and the rate limiting enzyme for TAG hydrolysis (15–17). These more recent data suggest that the I148M variant in PNPLA3 is a neomorph that confers a new function. Apart from PNPLA3’s role as a TAG lipase, it has also been shown to exhibit transacylase and acyltransferase activity, transferring long-chain fatty acid groups to acceptor substrates (i.e. neutral lipids) (13, 15, 18–20). Transacylase and acyltransferase reactions are known to be crucial in the formation and remodeling of cellular membranes as well as the storage of signaling lipids (21). However, WT PNPLA3 has relatively weak intrinsic enzymatic activity, suggesting it’s primary physiological role may revolve around interactions with protein binding partners such as ABHD5 (17).

MASLD is a multisystem disease which negatively impacts several extrahepatic organs, including the heart and vasculature (22). In fact, the primary cause of hospitalizations and mortality in MASLD patients is cardiovascular disease (CVD) (23). Risk of CVD in MASLD patients is most often attributed to increased levels of plasma lipids resulting from dysregulated hepatic lipid metabolism and increased secretion of very low density lipoprotein (VLDL) (24). Moreover, an updated meta-analysis of about 11 million individuals reports a role for MASLD in promoting new-onset heart failure through progression of cardiac arrhythmias and structural abnormalities (25). Currently, a mechanistic understanding for how MASLD promotes development of cardiac dysfunction is lacking.

Paradoxically, PNPLA3 I148M is associated with a reduced risk of CVD incidence and mortality in MASLD patients (26–28). Large scale genome- and exome-wide association studies report that human carriers of the GG allele (I148M) have lower levels of plasma lipids and a reduced risk of developing CVD (29–31). Indeed, several studies report involvement of PNPLA3 in the formation and secretion of VLDL particles (19, 32–34), suggesting a potential pathway for how PNPLA3 I148M might lead to protection against cardiovascular dysfunction. Here we employ a humanized mouse model of PNPLA3 expression, in which PNPLA3-/- mice with hepatic expression of human WT PNPLA3, human PNPLA3 I148M or a GFP control, were placed on a chow or MASH inducing diet to examine the effects of hepatic PNPLA3 expression on cardiovascular health in mice. We demonstrate that under a MASH diet, PNPLA3 I148M promotes fatty liver disease by reducing hepatic secretion of TAG rich particles, thereby likely eliciting a cardioprotective effect.

## METHODS

### Animal Studies

PNPLA3 knockout mice on a C57B6/J background were a generous gift from Dr. James Perfield (Eli Lilly) and were sourced from Taconic Biosciences (Cat# TF2065). Mice were bred and housed at standard room temperature with a 12 h light: 12 h dark cycle in an American Association for Laboratory Animal Care approved facility. All protocols involving animals were approved by the Institutional Animal Care and Use Committee at Wayne State University. Male mice between 8 and 12 weeks of age were tail vein injected with 1x10^6^ genome copies of adeno-associated virus-8 (AAV8) under a thyroglobulin promoter for hepatic specific expression of human wildtype PNPLA3, human PNPLA3 I148M or a GFP control. Mice were given 2 weeks to recover after injection with AAV, at which point they were switched from a standard chow diet to the Gubra Amylin NASH (GAN: 0910031040) diet, hereon referred to as Metabolic-Associated Steatohepatitis (MASH) diet (Supplemental Table 1).

Additionally, two weeks post AAV injections mice were moved from their ventilated cages kept at standard room temperatures (∼21-23°C) to thermoneutral housing between 29-30°C, by placing them in static cages in a Caron Scientific animal rearing chamber. Mice were maintained on either chow or MASH diet for 16 or 24 weeks at which point they were sacrificed using 3-5% isoflurane anesthesia and harvested for tissues.

### Indirect calorimetry

At ∼10 weeks of thermoneutrality indirect calorimetry was used to obtain measures of food intake, energy expenditure, respiratory exchange ratio and oxygen consumption by individually housing mice in a comprehensive animal monitoring system (CLAMS – Columbus Instruments). Housing temperatures were maintained at 29-30°C with a relative humidity between 20-40%. All mice were given ∼36-48 hours to acclimate to the chambers prior to collection of experimental data. For data collection air is sampled every 20 seconds from one of the housing units, which then repeats on a cycle for the duration of the collection period. Indirect calorimetry data was collected for at least 48-72 hours for all mice. All data measurements were analyzed using CLAMS examination tool software (CLAX) and exported to plot in GraphPad Prism.

### Gene expression qRT-PCR

Total RNA was isolated from RNA-Later preserved liver samples using Trizol homogenization and extraction method. Reverse transcription was completed using 1μg of total RNA per sample to obtain 1μg of cDNA for each qPCR reaction. Reactions were set up using SYBR green master mix and 50μg of purified cDNA combined with forward and reverse primers (Supplementary Table 1) for a total of 20μL reaction volumes run in a 3-step cycling protocol for 40 cycles; denature at 95°C for 5 sec, anneal at 60°C for 15 sec, extend at 70°C for 15 sec, with a pre-activation at 95°C for 2 min, using a QuantStudio 7. All values were normalized to housekeeping genes L27 and PPIA to wildtype controls using the delta delta ct methods (35).

### Western Blotting

Sodium Dodecyl Sulfate Polyacrylamide Gel Electrophoresis (SDS-PAGE) was performed under standard conditions using 10% pre-cast gels (Supplemental Table 1). Resolved proteins were transferred to polyvinylidene difluoride (PVDF) membrane paper following a brief 2 min activation in methanol. After proteins were transferred, the PVDF membranes were blocked for 1 hour at room temperature using 5% powdered skim milk in wash buffer. Primary PNPLA3 antibody (R&D systems, AF5208) incubations were completed at 4°C for 16 hours overnight, secondary incubations were completed at room temperature for 1 hour (anti-sheep diluted 1:20,000). Clarity Max ECL substrate (BioRad Cat. #1705062) was used for visualization of proteins.

### Hepatic Triacylglycerol Quantification

Neutral lipids were extracted from approximately 50mg piece of liver using the Folch method (36). Briefly, the liver piece was bead-homogenized in a precooled tube containing 1mL of 2:1 chloroform: methanol and ceramic beads followed by nutation at 4°C overnight.

The following day tubes were centrifuged at 7,000 rpm for 10 minutes at 4°C to pellet debris. The solution was transferred to a new tube and 200μL of sodium chloride solution was added to each sample and vortexed to ensure thorough mixing. Samples were left to stand at room temperature, allowing the organic and inorganic phases to separate. The organic supernatant (chloroform layer), was carefully transferred to a new glass tube and subsequently dried down using a constant stream of nitrogen gas, facilitating the removal of residual solvents. After drying, ACS-grade isopropanol containing 1% Triton X-100 was added followed by thorough vortexing to fully dissolve TAGs. Determination of hepatic TAG context was completed using a TAG determination kit (Sigma-Aldrich, TR0100) with the absorbance measured at 540nm using a BioTek Synergy plate reader. Data was normalized back to liver chip weight used for each sample for a comparative analysis.

### Echocardiography

To investigate cardiac function, echocardiography was performed ∼2 weeks post AAV injection, just prior to the start of MASH diet and thermoneutrality and then again at 16 weeks post diet at thermoneutrality. Internal body temperature was kept between 35-37°C while heart rate was maintained at 400-550bpm during all reported measurements (Figure 4B). Mice were anesthetized with isoflurane, 3% for induction and 1-2% for maintenance, under the control of a SomnoSuite low-flow anesthesia system. Hair covering the chest cavity was removed using non-scented Nair. Body temperature was recorded using a rectal probe and was maintained between 35-37°C when possible. Images of cardiac function were captured using Vevo3100 ultrasound, equipped with the MX400 transducer and analyzed by blinded investigator using VevoLAB analysis software. Two-dimensional images were captured using both parasternal long-axis and parasternal short axis views of the left ventricle for determination of ventricular wall thickness, mass and function. Determination of left ventricular early (E) to late (A) filling velocities (E/A ratios), which were used to assess diastolic function, were measured using pulse wave Power Doppler imaging and captured using the apical four-chamber view.

### Triacylglycerol secretion assay

Following a brief 4 hour fast, a 0 timepoint blood sample was collected from a nick in the tail vein followed by tail vein injected with tyloxapol (500mg/kg), an LPL inhibitor which prevents the peripheral uptake of VLDL-TAGs creating a gradual rise in plasma TAGs. Blood was then collected at 30, 60, 90 & 120 min from the time of injection in heparinized capillary tubes. Blood samples were centrifuged at 6,000 rpm for 10 min at 4°C to separate plasma which was collected and analyzed for TAGs using a kit (Sigma-Aldrich, TR0100). The same mice were then placed on a MASH diet for 4 weeks and the TAG secretion experiment was repeated exactly the same as described above.

### Plasma cholesterol assay

Plasma was isolated from final blood draws prior to sacrifice following 16 weeks of MASH or chow diet and a brief 4 hour fast. Plasma cholesterol levels were measured using a kit from Point Scientific (Point Scientific Ref. # C7510-120) as suggested by the manufacturer. Briefly, plasma samples were diluted 1:4 in 0.9% sterile saline and plated in duplicate. Cholesterol reagent was added to each well at which point plate was incubated for 5 min at 37°C prior to reading absorbance at 500nm.

### Intraperitoneal Glucose Tolerance & Insulin Sensitivity Tests

Following 12-13 weeks of thermoneutrality, mice were fasted at 6 am for 6 hours and all mice were subjected to a glucose tolerance test by way of a bolus intraperitoneal (IP) injection of glucose at a 0.1g/mL concentration for a final dose of 1g/kg. Blood was collected from a nick in the tail vein prior to glucose injection (time 0) and at 15, 30, 60, 90 & 120 min from time of injection and read on a glucometer (One Touch Verio). At 13-14 weeks of thermoneutrality, an insulin sensitivity test was performed where mice were given a bolus IP injection of insulin at a dose of 1 unit/mL/kg. Blood was collected as above and read on a glucometer.

### Quantification of atherosclerosis and Histology

Following 24 weeks of MASH diet, mouse hearts were dissected, formalin fixed and embedded in optimal temperature cutting compound (OCT) prior to freezing on dry ice. Hearts were sectioned down into the aortic root using a Leica cryostat at a thickness of 4-5μM. Sections were allowed to air dry prior to a brief wash in water and then 60% isopropanol. Following the washes slides were stained with a 0.1% working solution of Oil Red O (Ref. #O0625) for ∼5 minutes per slide. Slides were then washed again in isopropanol and water prior to staining with Harris’s Hematoxylin for about 30 seconds to provide a background to contrast the Oil Red O. After staining slides were mounted using an aqueous mounting medium and sealed using clear nail polish. Images were captured using a Leica Slide Scanner at 20X magnification and analyzed using ImageJ software. Blinded analysis of atherosclerotic plaque size was determined from 3-5 sections per mouse and area was calculated by manually outlining the plaques using the free hand tool in ImageJ. For histology, tissue pieces were fixed in 10% formalin, paraffin embedded and processed for hematoxylin & eosin staining.

### Lipidomics

To measure total liver, heart and plasma lipid composition, a total lipid extract was prepared using the method first published in Matyash *et al.,* 2008 (37). Briefly, tissues were snap frozen in liquid nitrogen and then pulverized into a fine powder using a supercooled Bessman Tissue Pulverizer and transferred to new tubes. Powdered samples were then shipped out on dry ice for liquid chromatography mass spectrometry (LC-MS) at the University of California Davis. A students t-test was performed comparing PNPLA3 I148M and WT PNPLA3 mice for the indicated lipids and the p-values were log transformed (-log(p)) and plotted against the log2 fold change (FC) between PNPLA3 I148M and WT PNPLA3. The values were plotted in a volcano plot format using GraphPad Prism software. Lipids that were significantly increased in PNPLA3 I148M were plotted in red, those that were decreased were plotted in blue and those that were not significant were plotted in grey.

### Statistical Analysis

All data are reported as mean ±SEM. Statistical significance was determined using GraphPad Prism software version 10. Data was analyzed with one-way analysis of variance (ANOVA), followed by Tukey’s post hoc, Two-way ANOVA, followed by Sidak’s multiple comparisons with and/or without repeated measures as indicated.

## RESULTS

### Hepatic expression of PNPLA3 I148M does not alter whole body metabolism

To understand how PNPLA3 I148M causes MASH and how it might protect against cardiac dysfunction, we utilized a humanized mouse model of PNPLA3 expression. Male PNPLA3-/- mice with hepatic specific expression of human wildtype (WT) PNPLA3, human PNPLA3 I148M or a GFP control were given a chow or MASH promoting diet for 16 weeks and housed at thermoneutrality (29-30°C). Thermoneutral housing has previously been shown to further exacerbate development of MASLD and promote induction of atherosclerosis in C57BL/6 mice fed a high fat or western diet (38–40). To assess the effects of hepatic PNPLA3 expression on whole body metabolism, indirect calorimetry was performed following 10-11 weeks of chow or MASH diet. No differences were found in food intake, energy expenditure, respiratory exchange ratios (RER), oxygen consumption, or bodyweights between the PNPLA3 groups fed a MASH diet (Figure 1A-E). At 12 weeks post MASH diet, mice expressing WT PNPLA3 demonstrated impaired glucose tolerance compared to GFP mice, while no difference was observed between mice expressing PNPLA3 I148M compared GFP and WT (Figure 1F). Insulin sensitivity was unaffected in mice following 12 weeks of MASH diet (Figure 1G). In mice fed a chow diet no changes in food intake, energy expenditure, respiratory exchange ratio, oxygen consumption, bodyweights, glucose tolerance, or insulin sensitivity were observed between the groups (Supplemental Figure 1).

**Figure 1:**
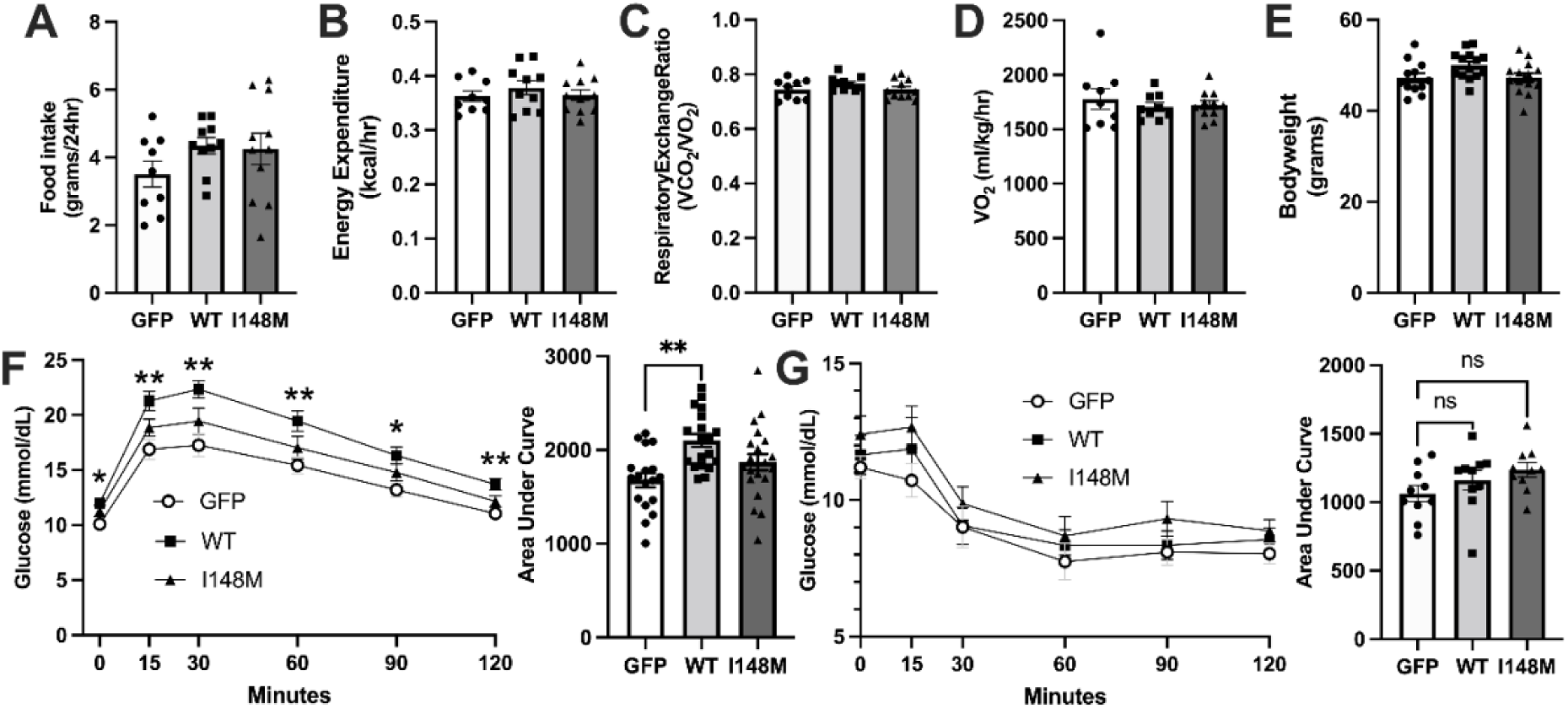
Whole animal metabolic phenotyping in mice following 12 weeks of MASH diet. Food intake (A), energy expenditure (B), respiratory exchange ratio (RER) (C), VO_2_ (oxygen uptake) (D), bodyweights at 11 weeks (E), intraperitoneal glucose tolerance (IP-GTT) at 12 weeks (F), and intraperitoneal insulin tolerance test at 13 weeks (IP-ITT) (G) in mice fed a MASH diet. Data represents mean +/- s.e.m. * *P* < 0.05 as indicated by one-way ANOVA (A-E) or * *P* < 0.05 and ** *P* < 0.01 by two-way ANOVA with repeated measures with a Tukey’s post-hoc analysis for glucose curves or one-way ANOVA for area under the curve analysis (F-G). Sample sizes were n= 9-12 mice per group except for glucose tolerance (n= 18-21 mice per group).

### Hepatic PNPLA3 I148M expression increases steatosis following MASH diet

Protein expression of human WT PNPLA3, PNPLA3 I148M and its absence in GFP mice was confirmed (Figure 2A). Following 16 weeks of MASH diet, mice expressing the I148M variant had greater liver to bodyweight ratios than did WT expressing mice (Figure 2B), while 16 week body weight and other tissue: body weights ratios were unchanged (Supplemental Figure 2A and B). No changes were detected in liver cholesterol after 16 weeks of MASH diet (Figure 2C). Notably, after 16 weeks of MASH diet all mice developed steatosis, with mice expressing PNPLA3 I148M showing a strong trend (p=0.071) for greater liver TAGs when compared to WT expressing mice (Figure 2D). Upon histological examination of H&E-stained liver sections, there appeared greater lipid vacuoles in mice expressing PNPLA3 I148M than either WT or GFP mice (Figure 2E). Steady state plasma TAG and cholesterol levels were unchanged between the groups following 16-weeks of MASH diet (Figure 2F-G). In chow fed mice, 16 week bodyweights and liver to bodyweight ratios were unchanged (Supplemental Figure 2C and 2D), while liver cholesterol was increased in chow fed I148M mice compared to GFP and WT, and liver TAGs were increased between WT and I148M (Supplemental Figure 2E and F). The increase in liver fat of I148M expressing mice did not result in any overt histological differences (Figure 2G). No differences were found in plasma TAGs or cholesterol in chow fed mice (Supplemental Figure 2 H-I). No differences were found in expression of inflammatory genes CCL2, CXCL-10, IL-6, IL-1β, from livers between the groups following 16 weeks of MASH diet at thermoneutrality, with the exception of GFP mice having greater TNF-α expression when compared to WT mice (Figure 3). Similarly, no changes were found in the expression of fibrosis related genes Acta2, Col1a1, MMP2, Timp1, TGF-β1, or TGF-β1 receptor between any of the groups following MASH diet (Figure 3).

**Figure 2:**
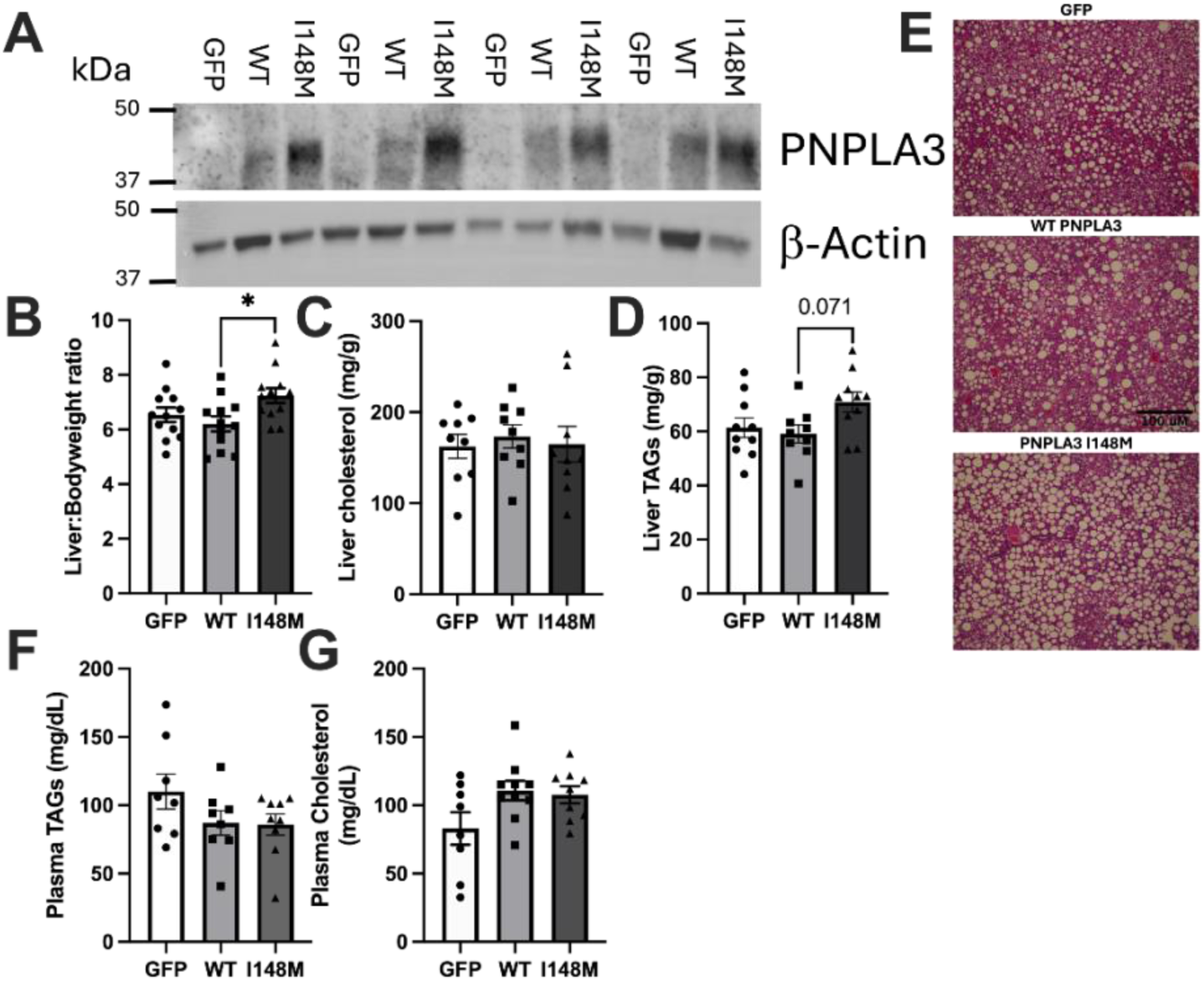
Biochemical analysis & histological examination of liver tissue following 16 weeks of MASH diet. Representative blot for human PNPLA3 in mouse liver tissue (A), liver to bodyweight ratios (B), liver cholesterol (C), liver triacylglycerols (D), representative hematoxylin & eosin stained liver sections (E), plasma TAGs, (F), plasma cholesterol (G). Data represents mean +/- s.e.m. * *P* < 0.05 as indicated by one-way ANOVA with Tukey’s post-hoc analysis. Sample sizes were n= 8-12 mice per group.

**Figure 3:**
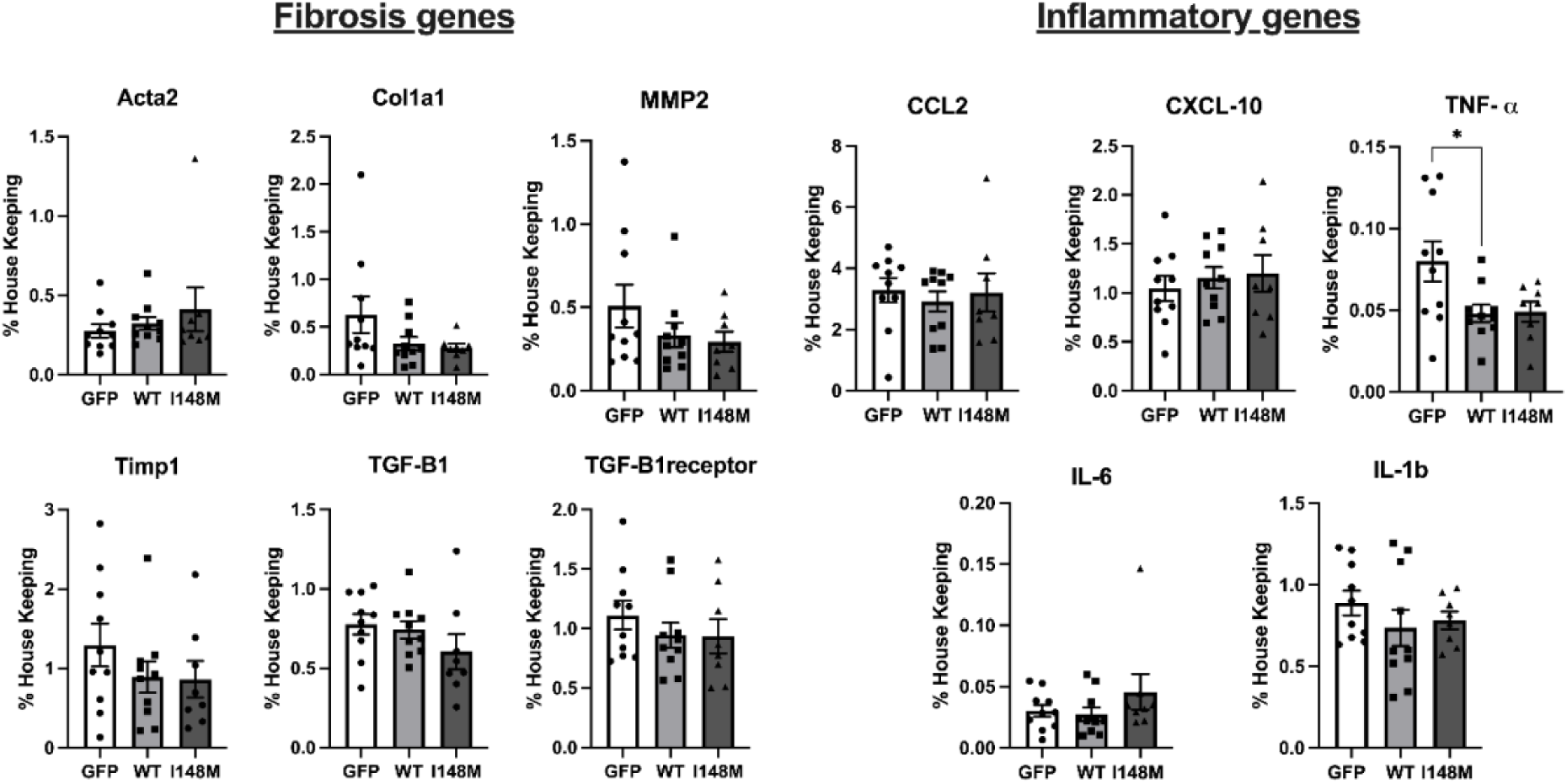
Hepatic fibrosis & inflammatory gene expression in mice fed a MASH diet for 16-weeks. Fibrosis genes (left): Acta2, Col1a1, MMP2, Timp1, TGF-β1, TGF-β1receptor. Inflammatory genes (right): CCL2, CXCl-10, TNF-α, IL-6, IL-1β. Data represents mean +/- s.e.m. * *P* < 0.05 as indicated by one-way ANOVA with Tukey’s post-hoc analysis. Sample sizes were n= 8-10 mice per group.

### Hepatic expression of PNPLA3 I148M protects against high-fat diet induced cardiac dysfunction in mice

To investigate cardiac functioning, echocardiography was performed 2 weeks post AAV injection, prior to the start of MASH diet and then again at 16 weeks post dietary intervention. Representative mitral flow Power Doppler echocardiography traces pre and post MASH diet for mice expressing liver GFP, WT PNPLA3, or PNPLA3 I148M are shown in Figure 4A. No differences were observed within any of the groups pre to post diet for heart rate, percent fractional shortening or percent ejection fraction, suggesting that systolic functioning was unaltered by 16 weeks of MASH diet at thermoneutrality (Figure 4B-D).

**Figure 4:**
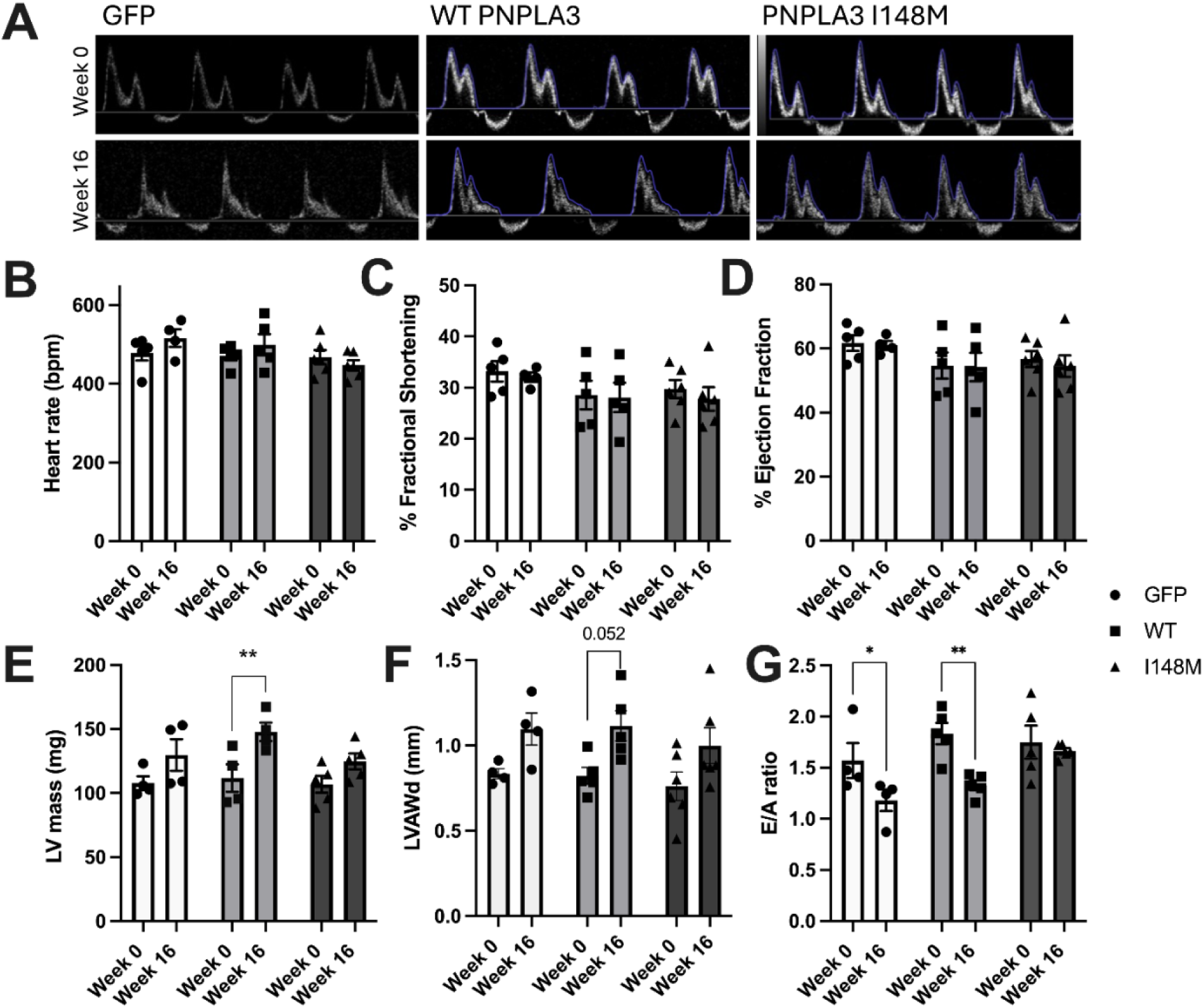
Echocardiography analysis of cardiac morphology and left ventricular diastolic function in mice following 16 weeks of MASH diet. Representative echocardiography traces of mice following 16 weeks of MASH Diet (A). Heart rates (beats per minute) at time of echocardiography (B), % fractional shortening (C), % ejection fraction (D), Left ventricular mass (E), Left ventricular anterior wall thickness during diastole (LVAWd) (F), Left ventricular relaxation E/A ratios (G). Data represents means +/- s.e.m. *p < 0.05 ** p < 0.01 as indicated by Two-way ANOVA with Tukey’s post-hoc analysis. Sample sizes were n= 4-6 mice per group.

Interestingly, mice with hepatic expression of WT PNPLA3 were found to have greater estimated left ventricular (LV) masses post MASH diet (Figure 4E), which was not detected in either the GFP or I148M mice. Along these same lines, WT PNPLA3 expressing mice displayed a strong trend (p=0.059) for increased left ventricle anterior wall thickness during diastole (LVAWd) post 16 weeks of MASH diet, which was not observed in either GFP or I148M expressing mice (Figure 4F). The E/A ratios, a measure of LV diastolic function, both GFP and WT PNPLA3 expressing groups displayed a reduction post MASH diet in E/A ratios which was not observed in the I148M group (Figure 4G). The preserved systolic measurements along with diet induced alterations in diastolic function suggest left ventricular diastolic dysfunction in mice expressing GFP and WT PNPLA3, in which PNPLA3 I148M mice seem to be protected from (Figure 4G). No changes were detected in cardiac function or morphology in age matched chow fed mice (Supplemental Figure 3), suggesting the effects on diastolic dysfunction were specific to the MASH diet.

### Hepatic expression of PNPLA3 I148M reduces VLDL-TAG secretion under MASH diet

To determine if the cardiac protection offered by PNPLA3 I148M was due to reduction in bulk secretion of lipids from the liver and whether this was dependent on diet, we examined liver TAG secretion in mice treated with the LPL inhibitor Tyloxapol, which blocks the peripheral uptake of TAGs from VLDL particles, enabling measurement of TAGs secreted from the liver. Under chow fed conditions, no differences in hepatic TAG secretion were observed between the groups (Figure 5A). However, when the same mice were fed a MASH inducing diet for 4 weeks, a significant reduction in hepatic TAG secretion was observed between mice expressing liver PNPLA3 I148M compared to WT PNPLA3 expressing mice and between PNPLA3 I148M and GFP at the indicated time points (Figure 5B) and an overall difference was observed between WT PNPLA3 and I148M.

**Figure 5:**
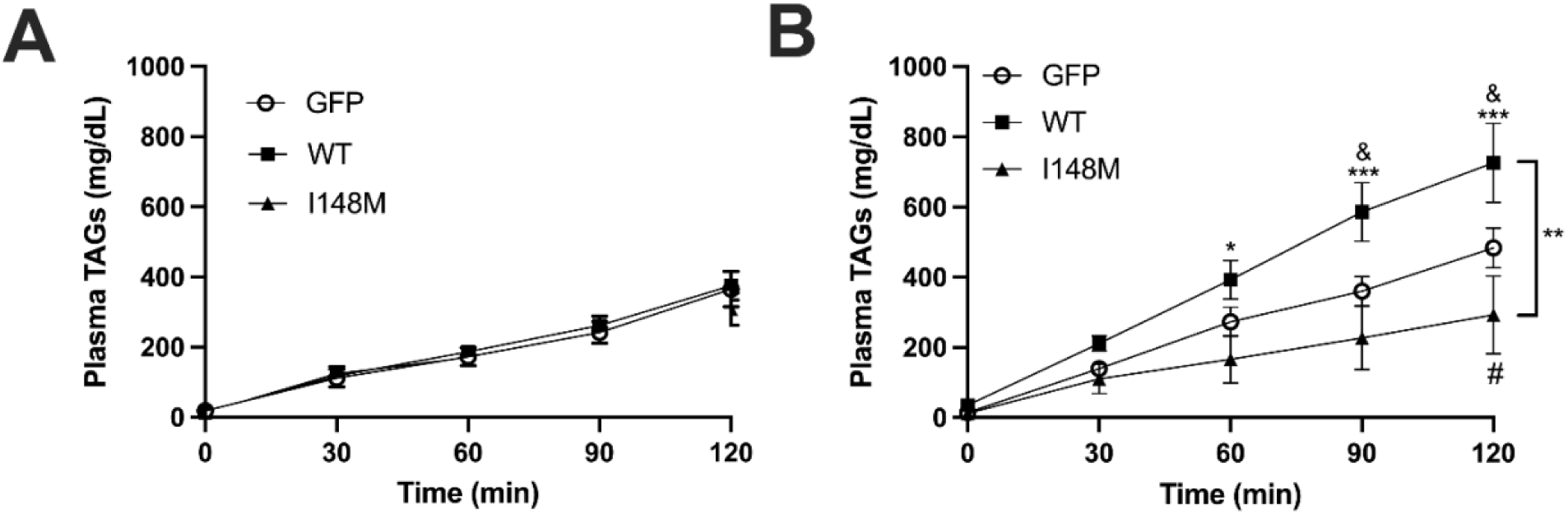
VLDL-TAG secretion in chow fed mice and following 4 weeks of MASH diet. Plasma TAG levels collected from chow fed mice following 500 mg/kg injection with Tyloxapol (A). Plasma TAG levels in mice fed four weeks of MASH diet following 500 mg/kg injection with Tyloxapol (B). Data represents means +/- s.e.m. * WT vs I148M, ^&^GFP vs WT, ^#^GFP vs I148M, *, ^&, #^ p < 0.05, *** p< 0.001 for indicated timepoints as indicated by two-way ANOVA with repeated measures (RM) and Holm-Sidak post-hoc analysis. An overall difference was observed between WT and I148M. ** p< 0.01 by one-way ANOVA with RM. Sample sizes were n= 5 mice per group for chow diet and n= 7-8 mice per group for MASH diet.

### Hepatic expression of PNPLA3 I148M alters lipidomes of liver and plasma with minimal changes in the heart

In addition to affecting TAG secretion, PNPLA3 I148M has been shown to modify the lipidome by promoting the sequestration of polyunsaturated fatty acids (PUFAs) in the liver, thereby preventing their incorporation into VLDL (19, 34). Moreover, omega-6 fatty acids are precursors to pro-inflammatory eicosanoids and the omega-6 PUFA arachidonic acid has been associated with cardiovascular dysfunction (41). Therefore, we examined if there were any changes in the liver lipidome and if this would result in complimentary changes to lipid species in the heart which might impact cardiac functioning. Unbiased lipidomic analysis of fatty acids, neutral lipids, phospholipids, ceramides sphingolipids and sterol lipids were performed in the liver, plasma and heart. In the liver, compared to WT PNPLA3, PNPLA3 I148M expression notably elevated the levels of several lipid species, including phosphatidylethanolamine 36:1, ether linked phosphatidylethanolamine PE O-18:1 20:3 and the omega-6 fatty acid docosadienoic acid 22:2 (Figure 6A). Other PUFAs such as 22:3, 24:3 and 24:4 approached significance (p=0.064; p=0.051 p=0.081 respectively). Conversely, significant decreases in PNPLA3 I148M liver compared to WT PNPLA3 were observed in various phosphatidylcholine (PC) and phosphatidylinositol (PI) species, notably PC-O 39:9 (ether linked phosphatidylcholine), PC 36:0, PC 22:4 and PI 40:6, among others (Figure 6A). Plasma lipidomic profiling revealed a significant reduction in 13-KODE (an octadecadienoic acid metabolite of linoleic acid), diacylglycerol 30:2, and myristic acid 14:0, alongside increases in phosphatidylglycerol 17:0 18:1, very long-chain polyunsaturated TAG 64:4, and ceramides 18:1 & 39:1 (Figure 6B). In contrast, minimal changes in heart lipids were observed between WT and PNPLA3 I148M with only a decrease in long-chain polyunsaturated TAG 68:2, sphingomyelin 40:0 and PC 33:2 levels, suggesting a minimal impact of liver PNPLA3 I148M on the heart lipidome (Figure 6C).

**Figure 6:**
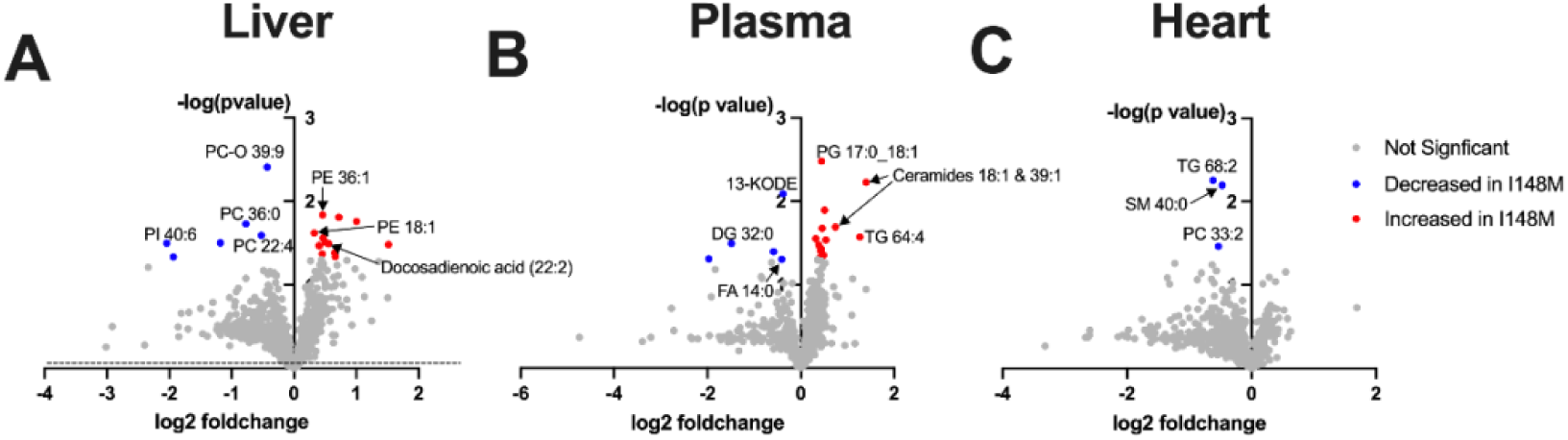
Untargeted complex lipidomic analysis of livers, plasma and hearts from 16 week MASH diet fed mice. Hepatic lipidome (A), Plasma lipidome (B), Cardiac lipidome (C). The -log (p-value) of PNPLA3 I148M vs WT was plotted against the log2 foldchange of the peak intensities for indicated lipid from liquid chromatography mass spectrometry data analysis. Comparison is between I148M and WT expressing mice. Sample sizes were n= 4-6 mice per group.

### Hepatic expression of PNPLA3 I148M does not alter atherosclerosis following 24 weeks of MASH diet

Carriers for PNPLA3 I148M are protected from CVD, specifically coronary artery disease. Given the reduction in hepatic TAG secretion observed with liver expression of PNPLA3 I148M under MASH-diet, we next evaluated atherosclerotic plaque formation. Mice were housed at thermoneutrality on a MASH diet for 24 weeks which has previously been shown to increase plaque formation (40). Following 24 weeks of MASH diet, plaque formation was observed (Figure 7A), however no differences in aortic ring size or size of the plaque region could be observed among the groups (Figure 7B and C).

**Figure 7:**
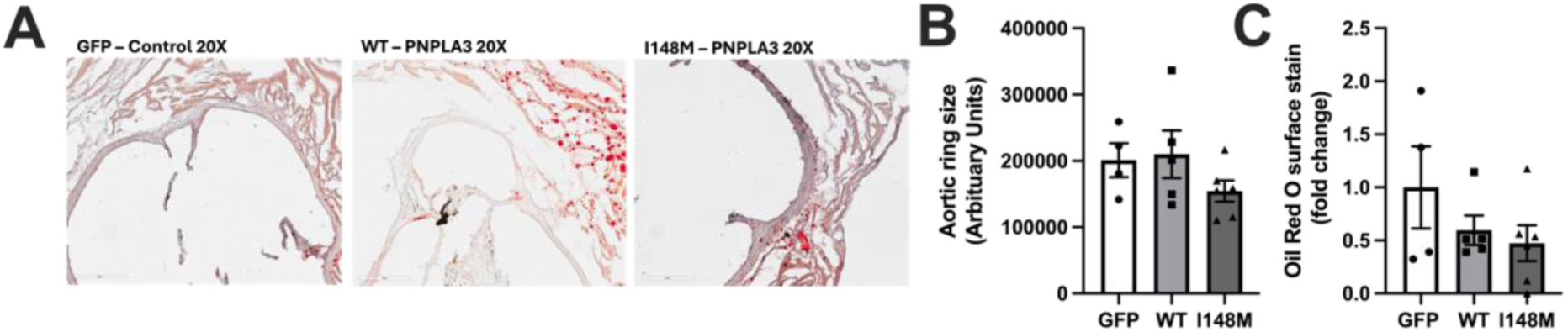
Quantification of atherosclerotic lesions following 24 weeks of MASH diet. Representative images of Oil-Red-O stained aortic roots for atherosclerotic lesion analysis (A). Aortic sinus lesion area (% of total area * 10^4^) (B), Total aortic sinus area (arbitrary units) (C). Data represents mean +/- s.e.m. Sample sizes were n= 4-6 mice per group.

## DISCUSSION

The PNPLA3 rs738409 variant, encoding the I148M substitution, has garnered significant attention as the greatest genetic risk factor for MASLD; however, interestingly this variant also appears to have a protective role against the development of cardiovascular disease in MASLD patients (26–30, 32, 42).

Here we investigated cardiovascular health in a humanized mouse liver model of PNPLA3 expression in mice fed a chow or MASH diet. We find that hepatic expression of PNPLA3 I148M preserves cardiac mass, thickness and diastolic function following 16 weeks of MASH diet, and this effect is likely due in part to PNPLA3 I148M reducing hepatic TAG secretion in MASH fed mice. We found that mice expressing human PNPLA3 I148M exhibited significant protection against diet-induced cardiac alterations, specifically increasing LV mass and decreasing E/A ratios, that was not observed in human WT PNPLA3 expressing counterparts (43). The unchanged LV mass and E/A ratios following 16 weeks of MASH diet in PNPLA3 I148M mice suggest a mitigated pathological response to metabolic stress, pointing to a cardioprotective mechanism associated with this mutation. Interestingly, no effect of chow diet was observed on cardiac function indicating the requirement of a dietary challenge to observe a cardioprotective effect of the I148M variant. These findings are particularly important for patients with MASLD who harbor the rs738409 mutation, as they suggest that the metabolic syndrome frequently seen in MASLD may not uniformly translate to cardiac dysfunction. For these individuals, the presence of PNPLA3 I148M could serve as a modifying factor influencing cardiovascular health, challenging the traditional view that MASLD invariably predisposes patients to heart disease.

PNPLA3 I148M has been suggested to promote hepatic TAG accumulation, in part by impairing hepatic secretion of TAG rich VLDL particles, consequently influencing systemic lipid profiles (27, 28, 33, 34). However, an effect of the I148M variant on lowering plasma lipids has not been consistently observed by all (12, 19) possibly due to differences in dietary factors. Here we examined the effects on TAG secretion in a controlled mouse model of humanized PNPLA3 expression and found that the I148M variant reduces liver TAG secretion under MASH diet conditions. PNPLA3 I148M resulted in greater accumulation of PUFA in the liver which was not observed in the heart, suggesting the mechanism by which the variant affects cardiac function is primarily through the bulk secretion of lipids. Such alterations in hepatic TAG secretion may lead to diminished accumulation of lipids within the epicardial adipose tissue or the myocardium itself, thereby preserving cardiac function (44). Obesity induced lipotoxicity of the heart is known to promote cardiac inflammation and fibrosis, impair ventricular relaxation and lead to mitochondrial dysfunction (45, 46).

Recent work demonstrates that patients with genetically linked MASLD display a lower relative risk of developing cardiovascular complications (47). Given its high populational prevalence, screening for the I148M variant will serve as an essential tool in disease risk stratification, allowing clinicians to identify high-risk individuals who may benefit from enhanced monitoring and preventative measures.

Furthermore, results from our studies suggest that MASLD patients who undergo PNPLA3 I148M knockdown or silencing therapies should be monitored for development of cardiovascular disease (48, 49). Overall, these results highlight the intricate relationship between hepatic lipid metabolism and cardiovascular health, warranting further exploration in genetically linked MASLD patient populations. In agreement with previous investigations, under chow fed conditions hepatic TAG secretion was unchanged by hepatic expression of WT PNPLA3 or I148M (12, 50). However, our data demonstrate that when fed a MASH inducing diet high in fructose, saturated fat and cholesterol, mice with hepatic expression of PNPLA3 I148M elicited a reduction in secreted TAG from the liver when compared to WT PNPLA3 expressing mice (Figure 5), suggesting that increased dietary fatty acid flux to the liver is required to observe an effect of PNPLA3 I148M on disrupting VLDL secretion. Surprisingly, in mice fed a fat-free high-sucrose diet, displayed no differences in VLDL-TAG secretion between mouse WT PNPLA3 and I148M, suggesting a dietary fat component that is required to reveal the effects of PNPLA3 I148M on TAG secretion or differences due to the mouse and human proteins (50). Recently Johnson *et al.,* found that under lipogenic conditions of a western style diet WT PNPLA3 facilitates hepatic VLDL-TAG secretion via polyunsaturated fatty acid incorporation into phospholipids (34). Interestingly, this function was found to be reduced by PNPLA3 I148M; however, the direct comparison of mice deficient in PNPLA3 to PNPLA3I148M was not performed. Our results suggest that under dietary conditions which promote obesity and MASLD, PNPLA3 facilitates hepatic TAG secretion, a function which is reduced by the I148M variant (17). Hepatic TAG secretion was different between WT and GFP mice at indicated timepoints, an effect lowered by PNPLA3 I148M compared to GFP, suggesting that the I148M variant may not be a simple loss of function mutation, but may be functioning in a dominant negative manner to suppress TAG secretion (Figure 5). Taken together, the effect of PNPLA3 I148M on reducing hepatic TAG secretion requires an increased nutritional load, likely due to the nutritional regulation of PNPLA3’s expression and stability, which is increased upon feeding and stabilization with lipid droplets (50–52). Furthermore, similar to how PNPLA3 I148M promotes hepatosteatosis through sequestration of ABHD5 Sherman *et al.,* reports that PNPLA3 I148M impairs secretion of Apolipoprotein B, a protein involved in the trafficking of VLDL-cholesterol, in an ABHD5 dependent manner (53). These findings suggest that PNPLA3 is a key player involved in regulating hepatic secretion of VLDL under excess dietary conditions and that the I148M mutant impairs it activity and disrupts ABHD5 function, a potential mechanistic link mediating its cardioprotective effects. A reduction in hepatic secretion of VLDL-TAGs by PNPLA3 I148M may lead to a reduction in lipid overload to the heart thereby providing protection against myocardial lipotoxicity and left ventricular diastolic dysfunction in MASLD patients (54), as increased storage of lipids within the heart have previously been linked to altered mitochondrial morphology and respiration (55).

WT PNPLA3 expression resulted in a modest yet significant reduction in glucose tolerance when compared to GFP expressing controls under conditions of a MASH diet. This observation aligns with previous investigations demonstrating that overexpression of PNPLA3 impairs glucose tolerance, while knockdown of PNPLA3 in leptin deficient db/db mice improves glucose tolerance (56). Similarly, PNPLA3 knockout mice fed high-fat diet for 15 weeks were found to have improved glucose tolerance when compared to wildtype littermates (14). Mechanistically, PNPLA3 may affect insulin secretion (57) and/or insulin induced Akt activation, either directly or indirectly through the production of ceramides, lipid mediators which are known to impair insulin signaling pathways (58). In contrast to previous studies, our investigation did not reveal variations in hepatic inflammatory gene expression between the PNPLA3 groups, suggesting that the presence of the I148M variant does not enhance hepatic inflammation or fibrosis in our model (59). It is possible that the pro-inflammatory and pro-fibrotic effects of the PNPLA3 I148M variant only become evident when it is expressed in other cell types, such as stellate and/or Kupffer cells (60). Assessment of atherosclerotic plaques did not yield any differences between PNPLA3 groups after 24 weeks on MASH diet. While mice are typically quite resistant to developing atherosclerosis, thermoneutral housing paired with a western diet has previously been shown to induce atherosclerotic lesion formation in laboratory mice (40). However, it is possible that the atherosclerosis observed in our mice fed a MASH diet for 24 weeks at thermoneutrality may not be enough to detect a protective effect by PNPLA3 I148M.

While this study provides insights into the cardioprotective role of PNPLA3 I148M in the context of MASLD, study limitations must be acknowledged. First, the use of murine models may not fully replicate human metabolic and cardiovascular responses, particularly interspecies differences in lipid metabolism and disease pathogenesis. Moreover, this study utilized male mice which may limit the generalizability of the findings to female populations where hormonal differences could influence disease progression (61). Additionally, while the humanized mouse model of PNPLA3 expression illustrates a role for PNPLA3 in hepatic TAG secretion and diet induced cardiac protection, it does not account for the multifactorial nature of MASLD and cardiovascular disease development, including interactions with other genes, environmental pollutants and/or additional dietary considerations such as alcohol consumption.

In conclusion, this study highlights the role of PNPLA3 I148M in promoting MASLD by reducing hepatic TAG secretion and mitigating diet induced cardiac alterations associated with metabolic dysregulation in MASH diet fed mice. Using a humanized mouse model, we demonstrate that hepatic expression of PNPLA3 I148M preserves cardiac structure and diastolic function, likely due to its impact on reducing hepatic TAG secretion in response to a MASH diet. Notably, these findings challenge the conventional paradigm that MASLD invariably predisposes to cardiovascular complications, suggesting that the presence of the I148M variant may confer protective benefits for cardiac health in MASLD patients. The alterations in hepatic lipid metabolism and the systemic implications on cardiac health underscore the need for personalized approaches in treating patients with MASLD. Future investigations should focus on elucidating the underlying molecular mechanisms for how PNPLA3 I148M decreases hepatic TAG secretion under conditions of a MASH diet and how this translates to cardiac protection.

## DATA AVAILABILITY

All data reported in this paper will be shared by the corresponding contact.

## Author Contributions

E.P.M. conceived the experiment, E.P.M., A.J.B., J.G.G. designed the experiments A.J.B., E.P.M., N.T., G.T., A.A.R., K.A.A conducted the experiments. A.J.B., E.P.M., P.K., analyzed the data and A.J.B., and E.P.M. wrote and edited the manuscript. All authors read the manuscript and approved the final version.

## Supporting Information

This article contain supplementary materials.

## Acknowledgements

We thank Dr. James Perfield (Eli Lilly and Company) for providing AAVs and access to the PNPLA3-KO mice.

## Abbreviations

AAV: adeno-associated virus
ABHD5: alpha/beta hydrolase domain-containing protein 5
CLAMs: comprehensive animal monitoring system
CVD: cardiovascular disease
GAN: Gubra amylin NASH diet
LV: left ventricular
MASLD: metabolic dysfunction-associated steatotic liver disease
PNPLA2: Patatin Like Phospholipase Domain Containing 2
PNPLA3: Patatin Like Phospholipase Domain Containing 3
PVDF: polyvinylidene difluoride
SDS-PAGE: Sodium Dodecyl Sulfate Polyacrylamide Gel Electrophoresis
TAG: Triacylglycerol; WT, wildtype

