## Supplemental Data for "PNPLA3 I148M Reduces Hepatic Triacylglycerol Secretion and Mitigates Left Ventricular Diastolic Dysfunction in MASH Diet Mice"

**SUPPLEMENTAL INFORMATION**

**Supplementary Figure 1**

**Supplementary Figure 2**

**Supplementary Figure 3**

**Supplementary Table S1**

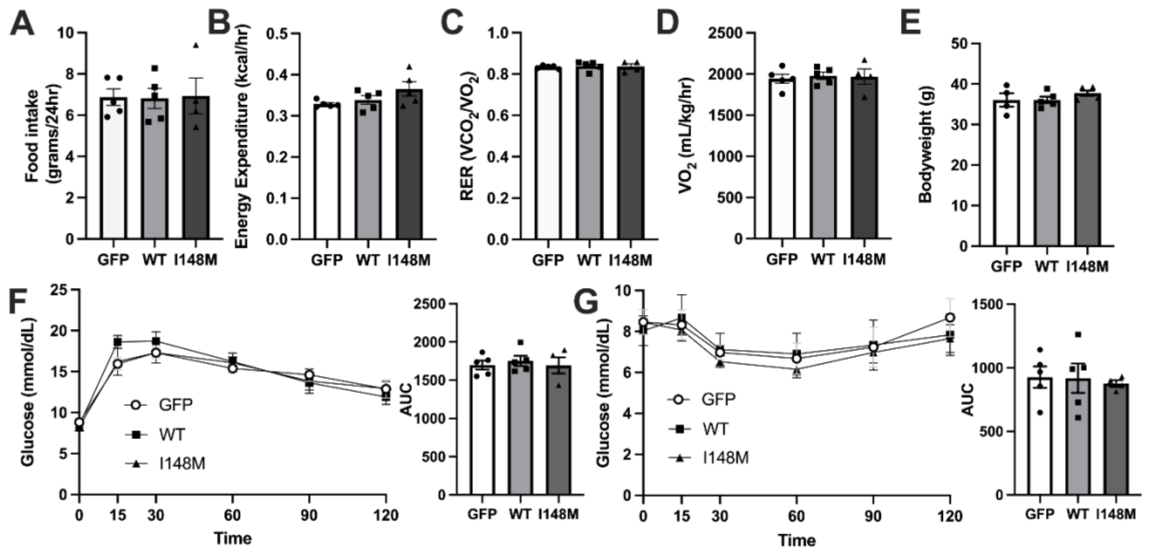


**Supplemental Figure 1:** Whole body metabolic phenotyping in age matched chow fed mice. Food intake (A), Energy expenditure (B), Respiratory exchange ratio (RER) (C), $VO_2$ (oxygen uptake) (D), Bodyweights at time of CLAMS (E), intraperitoneal glucose tolerance test (F), and intraperitoneal insulin tolerance test (G). Data represents mean +/- s.e.m. Data analyzed by one-way ANOVA or two-way ANOVA with a Tukey's post-hoc analysis. Sample sizes were n= 4-5 mice per group.

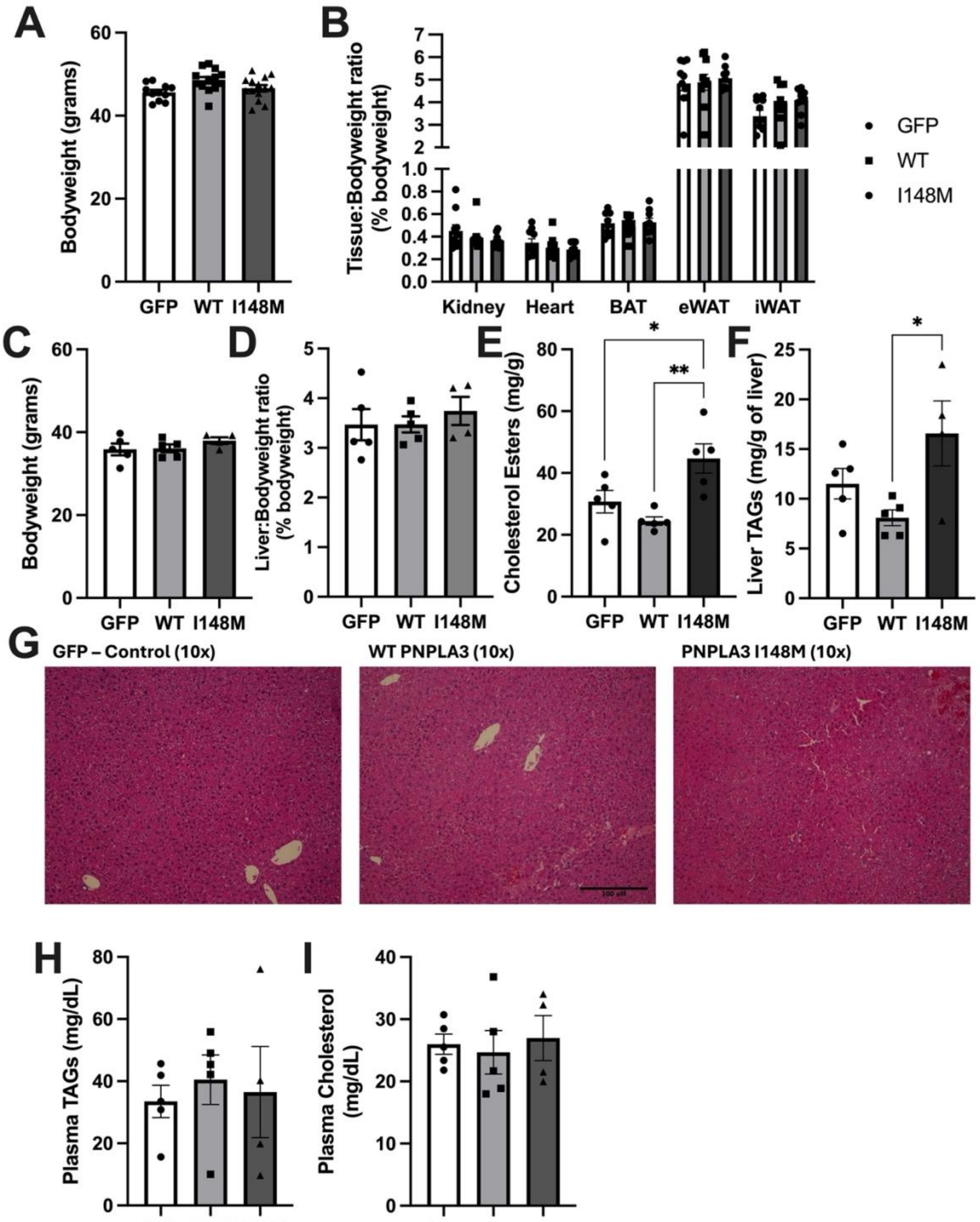

A
B
C
D
E
F
G
H
I
Bodyweight (grams)
GFP
WT
I148M
Tissue:Bodyweight ratio (% bodyweight)
Kidney
Heart
BAT
eWAT
iWAT
Liver:Bodyweight ratio (% bodyweight)
Cholesterol Esters (mg/g)
Liver TAGs (mg/g of liver)
GFP – Control (10x)
WT PNPLA3 (10x)
PNPLA3 I148M (10x)
Plasma TAGs (mg/dL)
Plasma Cholesterol (mg/dL)

**Supplemental Figure 2:** Biochemical analysis & histological examination of liver tissue from age matched chow fed mice. Bodyweights 16 weeks post MASH diet (A), Tissue to bodyweight ratios from 16 week MASH diet fed mice (B), bodyweights post 16 weeks chow diet (C), liver to bodyweight ratios (D), liver cholesterol (E), liver triacylglycerols (F), representative H&E-stained liver sections (G), plasma TAGs (H), and plasma cholesterol (I) in age matched chow fed mice at 16 weeks. Data represents mean +/- s.e.m. Statistical significance between genotypes was determined by a one-way ANOVA with Tukey's post-hoc; $*p < 0.05$ $**p < 0.01$. Sample sizes were n= 4-5 mice per group, except A-B n= 9-12.

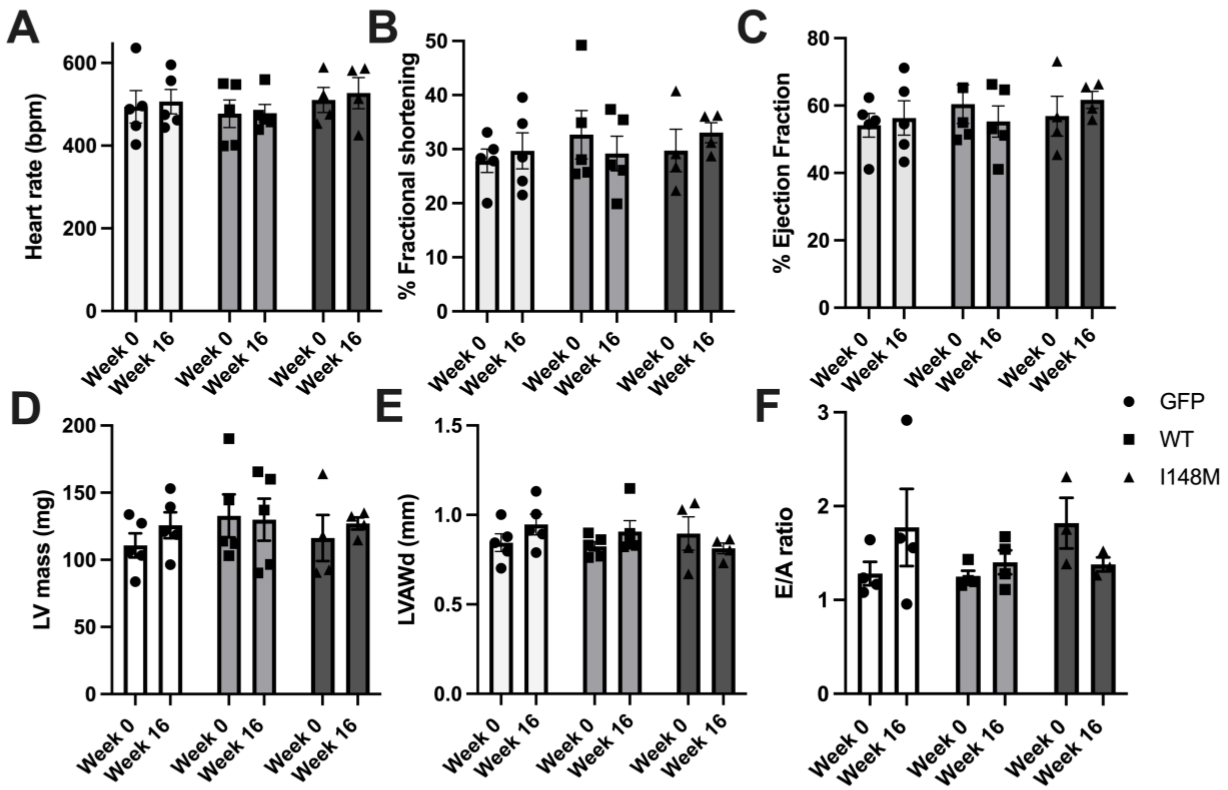


**Supplemental Figure 3:** Echocardiography analysis of cardiac morphology and left ventricular diastolic function in age matched chow fed mice two weeks following AAV injection (week 0) and following 16 weeks of chow (week 16). Heart rate (beats per minute) during echocardiography (A), Percent fractional shortening (B), Percent ejection fraction (C), Left ventricular (LV) mass (D), Left ventricle anterior wall thickness during diastole (LVAWd) (E), E/A ratio of the left ventricle (F). Data represents means +/- s.e.m and was analyzed by a 2way ANOVA with repeated measures with a Tukey's post-hoc analysis (A-F). Sample sizes were n= 4-5 mice per group.

**Supplementary Table 1: List of reagents**

| Reagent Type | Designation | Source | Additional information |
|---|---|---|---|
| Primer | IL-1b | Sigma | F: GGACCTTCCAGGATGAGGACA<br>R: GTTCATCTCGGAGCCTGTAGTG |
| Primer | IL-6 | Sigma | F: AGTGGCTAAGGACCAAGACC<br>R:TCTGACCACAGTGAGGAATG |
| Primer | Cxcl10 | Sigma | F: CCAAGTGCTGCCGTCATTTTC<br>R: GGCTCGCAGGGATGATTTCAA |
| Primer | TNF-a | Sigma | F: GGTGCCTATGTCTCAGCCTCTT<br>R: GCCATAGAACTGATGAGAGGGAG |
| Primer | Col1a1 | Sigma | F: GCTCCTCTTAGGGGCCACT<br>R: CCACGTCTCACCATTGGGG |
| Primer | MMP-2 | Sigma | F: CAAGTTCCCCGGCGATGTC<br>R: TTCTGGTCAAGGTCACCTGTC |
| Primer | TIMP1 | Sigma | F: GCAACTCGGACCTGGTCATAA<br>R: CGGCCCGTGATGAGAAACT |
| Primer | TGF-b1 | Sigma | F: CTCCCGTGGCTTCTAGTGC<br>R: GCCTTAGTTTGGACAGGATCTG |
| Primer | TGF-b1 receptor | Sigma | F: TCTGCATTGCACTTATGCTGA<br>R: AAAGGGCGATCTAGTGATGGA |
| Primer | Acta2 | Sigma | F: TGCTGACAGAGGCACCACTGAA<br>R: CAGTTGTACGTCCAGAGGCATAG |
| Antibody | Anti-sheep IgG PNPLA3 | R&D systems | Cat #: AF5208<br>Diluted 1:1,000 |
| Commercial assay or kit | TAG kit | Sigma-Aldrich | TR0100 |
| Commercial assay or kit | Total Cholesterol | Point Scientific | REF C7510-120 |
| Commercial assay or kit | AST kit | eLab Sciences | Cat #: E-BC-K236-M |
| Chemical compound, drug | Tyloxapol | Sigma-Aldrich | 500 mg/kg |
| Chemical compound, drug | Oil Red O | Sigma-Aldrich | |
| Rodent chow diet | Chow Diet 5L0D | PicoLab | |
| Rodent MASH diet | Gubra-amylin-NASH diet | Research diets | D#:0910031040% kcal fat (palm oil) 20% kcal fructose 2% cholesterol |
